# Resource asynchrony and landscape homogenization as drivers of virulence evolution

**DOI:** 10.1101/2022.08.03.502590

**Authors:** Tobias Kürschner, Cédric Scherer, Viktoriia Radchuk, Niels Blaum, Stephanie Kramer-Schadt

## Abstract

In the last years, the emergence of zoonotic diseases and the frequency of disease outbreaks have increased substantially, fuelled by habitat encroachment and asynchrony of biological cycles due to global change. The virulence of these diseases is a key aspect for their success. In order to understand the complex processes of pathogen virulence evolution in the global change context, we adapted an established individual-based model of host-pathogen dynamics. Our model simulates a population of social hosts affected by an evolving pathogen in a dynamic landscape. Pathogen virulence evolution is explored by the inclusion of multiple strains in the model that differ in their transmission capability and lethality. Simultaneously, the host’s resource landscape is subjected to spatial and temporal dynamics, emulating effects of global change.

We found an increase in pathogenic virulence and a shift in strain dominance with increasing landscape homogenisation. Our model further shows a trend to lower virulence pathogens being dominant in fragmented landscapes, although pulses of highly virulent strains are expected under resource asynchrony. While all landscape scenarios favour coexistence of low and high virulent strains, when host density increases, the high virulence strains capitalize on the high possibility for transmission and are likely to become dominant.

**Author Summary:** Disease outbreaks primarily caused by contact with animals are increasing in recent years, related to habitat destruction and altered biological cycles due to climate change. Pathogens associated with such outbreaks will be more successful the more effectively they can spread in a population. Thus, understanding the conditions over which those pathogens evolve will help us to limit the impact of disease outbreaks in the future. To this end, we used an individual based model that allowed us to study different scenarios. Our model had three main components: a host-pathogen system, a dynamic resource landscape with different degrees of fragmentation and temporal resource mismatches. We used dynamic landscapes with varying resource amounts over the years and consisting of multiple large or smaller habitat clusters. Our simulations showed that homogenous landscapes resulted in higher virulent pathogens and fragmented landscapes in lesser virulent pathogens. However, across all scenarios, high and low virulent pathogen strains were able to coexist.

## Introduction

A key aspect of the invasive success of infectious pathogens such as Ebola, SARS-CoV-2 or Avian Influenza in a host population is the mastering of the delicate interplay of transmission and host exploitation, also termed virulence. To persist, a pathogen must find the balance between quick replication and growth in the host often resulting in severe infections killing its host while still being able to spread across timescales (Visher et al. 2021). This intricate balance can only be kept up by an arms race between hosts’ immune reactions and strategies of the pathogen to evade and counteract host resistance, termed adaptive evolution of virulence (Cressler et al. 2016). This leads to the emergence of ever-new pathogenic strains from the wild strain with modulated pathogenic traits, and if the new strain manages to establish, this might have unforeseeable effects on host population and disease dynamics.

Both transmission and virulence are integrally tied to density and spatiotemporal distribution of host individuals (Alizon et al. 2009, Cressler et al. 2016), which in return are subject to habitat configuration and spatiotemporal variation in resource availability.

Global change might exacerbate disease dynamics in the near future, facilitated by land-use change, habitat encroachment, or climate warming (Patz et al. 2004, Wilcox and Gubler 2005). These disturbances will severely influence disease outbreaks by changes in the life history, density and availability of hosts as well as feedbacks on the landscape level due to asynchrony in timescales (Kürschner et al. 2021). In this context, it is particularly important to not only understand factors that govern the spread and the persistence of pathogens in changing landscapes to put counteractive measures in place (Griette et al. 2015), but to also understand how these factors reciprocally influence the adaptive potential of pathogenic traits.

Virulence evolution is often highly accelerated during the emergence or invasion stage of an epidemic (Griette et al. 2015, Geoghegan and Holmes 2018). The emergence stage is characterized by a high number of susceptible —and later infected— host individuals associated with a high number of mutations due to the steep increase of infected individuals (Galvani 2003). Since the distribution of host individuals in a landscape determines the number of available susceptible individuals, local and regional host densities are important factors in the evolution of virulence (Boots 2004). With global change further altering the resource distribution in space and time, subsequent changes in the spatiotemporal density and distribution of host individuals (Galvani 2003, Boots 2004, Geoghegan and Holmes 2018) could influence the evolution of virulence. Density changes could for example be induced via mismatches between the host’s life history such as reproduction and host resource availability at that time.

Theory predicts an evolution towards low virulence through altered habitat configuration or host density distribution (Boots and Mealor 2007, Cressler et al. 2016). Virulence has been shown to be adaptive if there is a correlation with other pathogenic traits such as contagiousness, which is known as virulence-transmission trade-off hypothesis (Day 2003). The transmission-virulence trade-off hypothesis states that an increase in strain transmission causes shorter infections through higher lethality (Anderson and May 1982, Alizon and Michalakis 2015). In other words, pathogen virulence is subject to a variety of evolutionary trade-offs (Kamo et al. 2007, Messinger and Ostling 2009, Cressler et al. 2016).

The theoretical models of virulence evolution, particularly the classical adaptive dynamics framework, rely on the assumption that mutation of pathogens happens very slowly and that mutations towards new strains can only occur after the dominant strain has reached equilibrium (Dieckmann et al. 2005). However, such simplified assumptions are rarely applicable to pathogens in nature, which often undergo transient dynamics, for example due to temporal and spatial changes in the landscape structure. Due to temporal variation in the landscape, the formation of spatial (figure 1 a) and or temporal (figure 1 b) host niches can cascade through the density distribution of potential hosts onto host-pathogen interactions (figure 1 c, d). The formation of niches with varying beneficial or detrimental properties for host and pathogen could facilitate the appearance of different pathogenic strains at specific times or locations. The result can be a complex system of different competing and coexisting pathogen strains (figure 1 e) with their own spatial and temporal dynamics. The constant emergence, re-emergence, and extinction of pathogenic strains will result in an overlap and possible coexistence between different strains, all competing for the same resource (Choua and Bonachela 2019).

**Figure 1:**
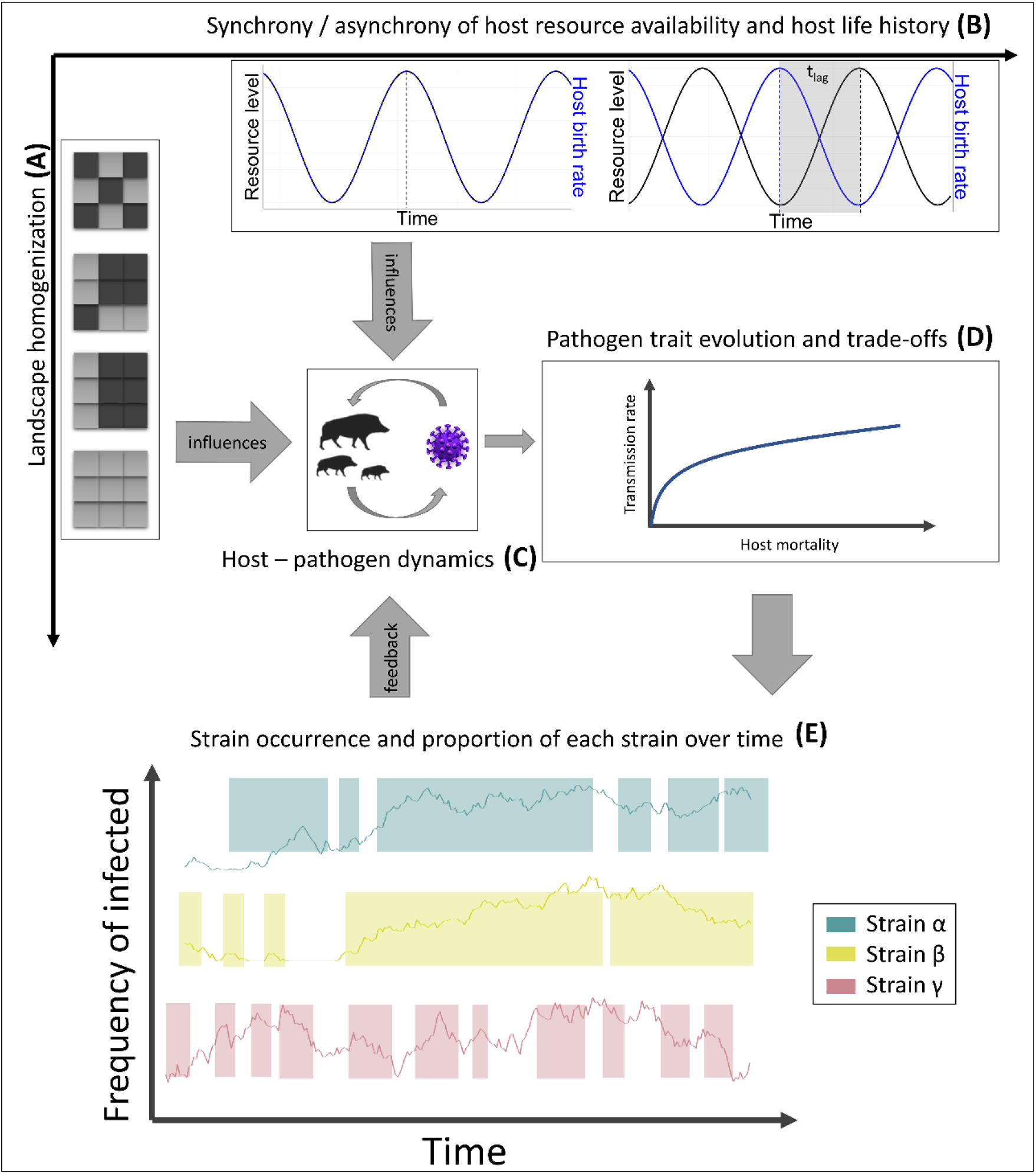
Conceptual figure: Landscape homogenization (A) and synchrony/asynchrony (t_lag_) of host life-history and host-resource availability (B) influence host-pathogen dynamics (C) and subsequently the evolution of pathogenic traits (D) that will affect strain occurrence over time where gaps in the background line are times when the strain did not occur in the landscape (E).

While theoretical studies focus on long-term predictions of pathogenic strains with evolutionarily stable virulence at equilibrium (Lenski and May 1994, Day and Gandon 2007), there is a lack of knowledge linking complex dynamics arising from global change to the evolution of virulence through space and time during an epidemic (Lebarbenchon et al. 2008). Also, links between resources and host density are rarely incorporated into evolutionary models, which typically assume that host density remains at equilibrium [3,28 in Hite&Cressler2018]. A prominent example tackling the evolution of virulence in changing host densities due to changes in resources is the work of Hite & Cressler (2018). They revealed complex effects of host population dynamics on parasite evolution, including regions of evolutionary bistability, where parasites ‘rode the cycles’ of their hosts and phases with high host exploitation superseded phases of low virulence (Hite & Cressler 2018).

Here, we go one step beyond the important link between host ecology and parasite evolution by asking what effect heterogeneously distributed and dynamic resources will pose on the evolution of virulence, particularly how temporal mismatches between optimal resource availability and biological events, such as reproduction, affect host-pathogen coexistence and pathogen spread through adaptive virulence dynamics. To this end, we modified an existing spatially-explicit individual-based host-pathogen model of a group-living social herbivore (Kramer-Schadt et al. 2009, Lange et al. 2012a, b, Scherer et al. 2020, Kürschner et al. 2021) and added evolution in pathogen traits leading to multi-strain outbreak scenarios. In accordance with theory, we have already shown for a static host exploitation rate that pathogen extinction is higher in landscapes with randomly distributed and fluctuating resources, but that the formation of disease hotspots form an epidemic rescue for the pathogen when hosts are mobile (Kürschner et al. 2021).

We here hypothesized that dynamic landscapes induce evolution in pathogenic virulence to facilitate host-pathogen coexistence (H1). In more detail, we expect pathogenic virulence to evolve into a system of different viral strains that will coexist and persist within the host population in parallel (prediction 1). We also predict that the frequency of ‘host cycle riding’ pathogenic strain emergence will be larger under environmental uncertainty, hence global change effects might lead to higher pathogenic strain emergence (prediction 2), i.e. with a higher chance for spill-over events.

We further hypothesize that due to the destabilisation of the host population under asynchronous dynamics, virulence will evolve to lower levels than under homogeneous and stable resource availability (H2). We expect increasing landscape homogenization and related contact homogenization to facilitate evolution towards higher pathogenic virulence by increasing the availability of hosts for highly virulent strains (prediction 3), with few dominant strains governing the dynamics for a long time (prediction 4).

## Results

### Host-pathogen coexistence

Overall, host pathogen coexistence P_coex_ was very high in almost all tested scenarios. Due to a collapse of the host population in asynchronous scenarios in homogeneous landscapes, host-pathogen coexistence was not achievable in the current model-framework and this single scenario therefore excluded. We did not find notable differences in the other scenarios (Appendix Fig. B2).

### Categorized infection trends

Our model showed that in synchronous scenarios, highly virulent strains were the least abundant ones among the three strain categories during the early stages of the epidemic. However, these strains became dominant in the later stages of the epidemic in homogeneous and large clustered landscapes (figure 2, left). With increasing landscape homogenization, medium virulence strains in the later stages of the epidemic were usually dominating along with high virulence strains. Across all landscapes, low virulent strains only occurred in high prevalence in the early stages of the epidemic but reached higher prevalence in less heterogeneous landscapes.

**Figure 2:**
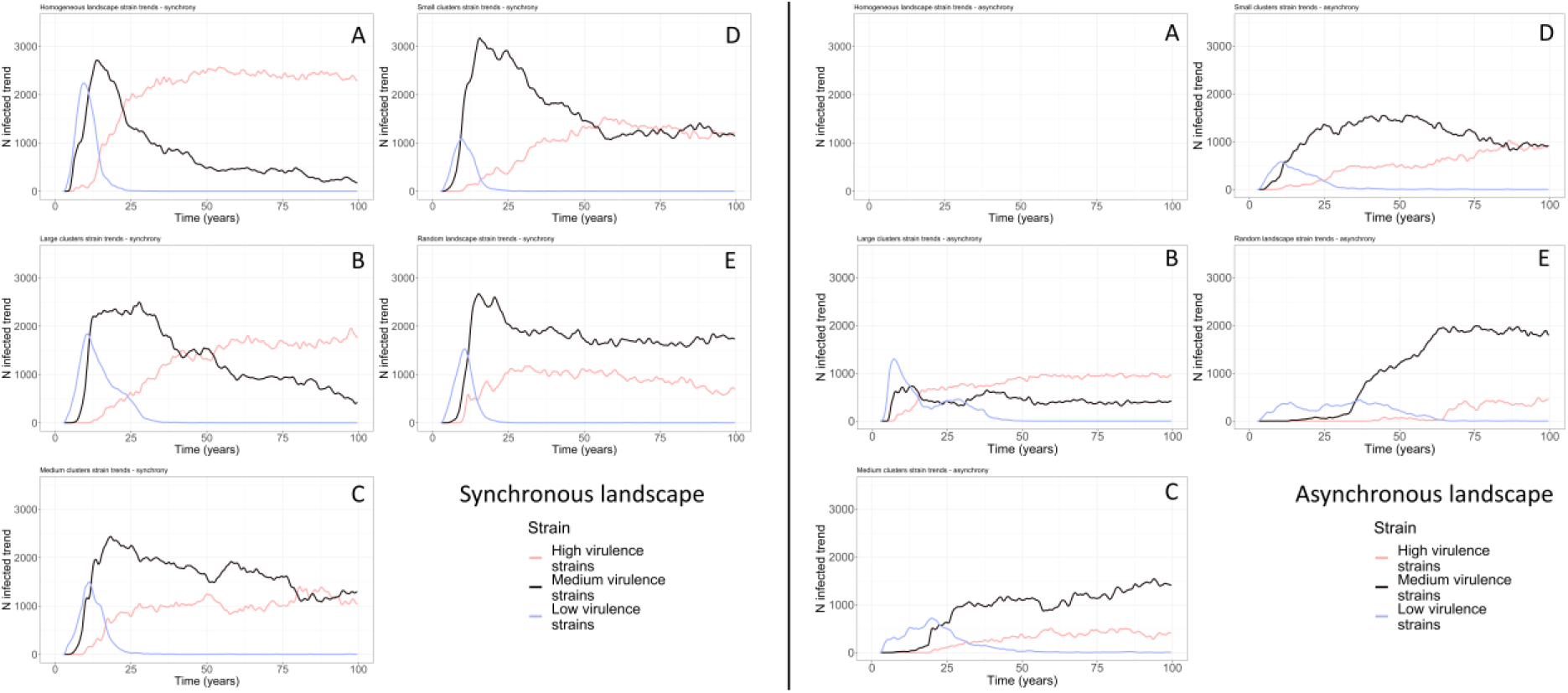
Temporal trends of the number of hosts infected with the strains of three virulence categories, low (blue) medium (black) and high (red) virulence over time. The left half shows the trends for synchronous host reproduction (t_lag_ = 0) and the right half for asynchronous host reproduction (t_lag_ = 100) scenarios.

In scenarios with asynchrony, low virulent strains occurred over a longer time period and were more prevalent in the host population, while medium and highly virulent strains occurred later at high prevalence (figure 2, right). Furthermore, prevalence of all strain categories was lower throughout the simulations when directly compared to the ‘synchronous’ scenarios. A clear shift towards a dominance of highly virulent strains only occurred in the less heterogeneous, large clustered landscapes.

#### Strain specific occurrence and dominance over time

Strain occurrence and dominance of high virulent strains decreased with increasing landscape heterogenization in dynamic landscapes in combination with synchronous scenarios (figure 3 top). In the highly heterogeneous random landscape, low virulent strains persisted longer while higher virulent strains occurred much later in time (around year 40), compared to the same landscape in synchronous scenarios. After its appearance through mutations around year 45, the medium virulent strain 7 became dominant for the course of the epidemic. As landscape homogeneity increased (from random landscapes to small clustered landscapes), lower virulent strains occurred on much shorter time spans. The high virulent strains appeared much earlier around year 10 of the epidemic and remained present in the landscape for the duration of the simulation. A further increase in homogenization towards medium-sized patches showed overall similar patterns as the small clustered landscape with the exception of the strain 7 peak prevalence, which shifted towards the end of the epidemic. In the highly homogenous large clustered landscapes, the lower virulent strains persisted for a longer period, while medium virulent strains were represented during the full period of the epidemic. Contrary to the more heterogeneous landscapes, strain 7 did not become the dominant strain despite high prevalence in the host population. As indicated by the larger proportion of hosts infected with higher virulent strains, overall, in synchronous scenarios (figure 3), virulence of occurring strains increased over time and with increasing landscape homogenization. In asynchronous scenarios we observed a similar increase in strain occurrence with landscape homogenization, even though the temporal rate of increase was smaller compared to synchronous scenarios. Furthermore, there was a temporal variation of strain occurrence within the more homogenous landscape between synchronous and asynchronous scenarios, as can be seen in a direct comparison of the per-strain infection counts and wavelets of synchronous and asynchronous scenarios over time (figure 4). A general comparison of the proportional strain contribution in asynchronous vs synchronous scenarios further showed that the occurring strains were of lower virulence in asynchronous scenarios in case of the more heterogeneous landscapes (Appendix Fig.B3).

**Figure 3:**
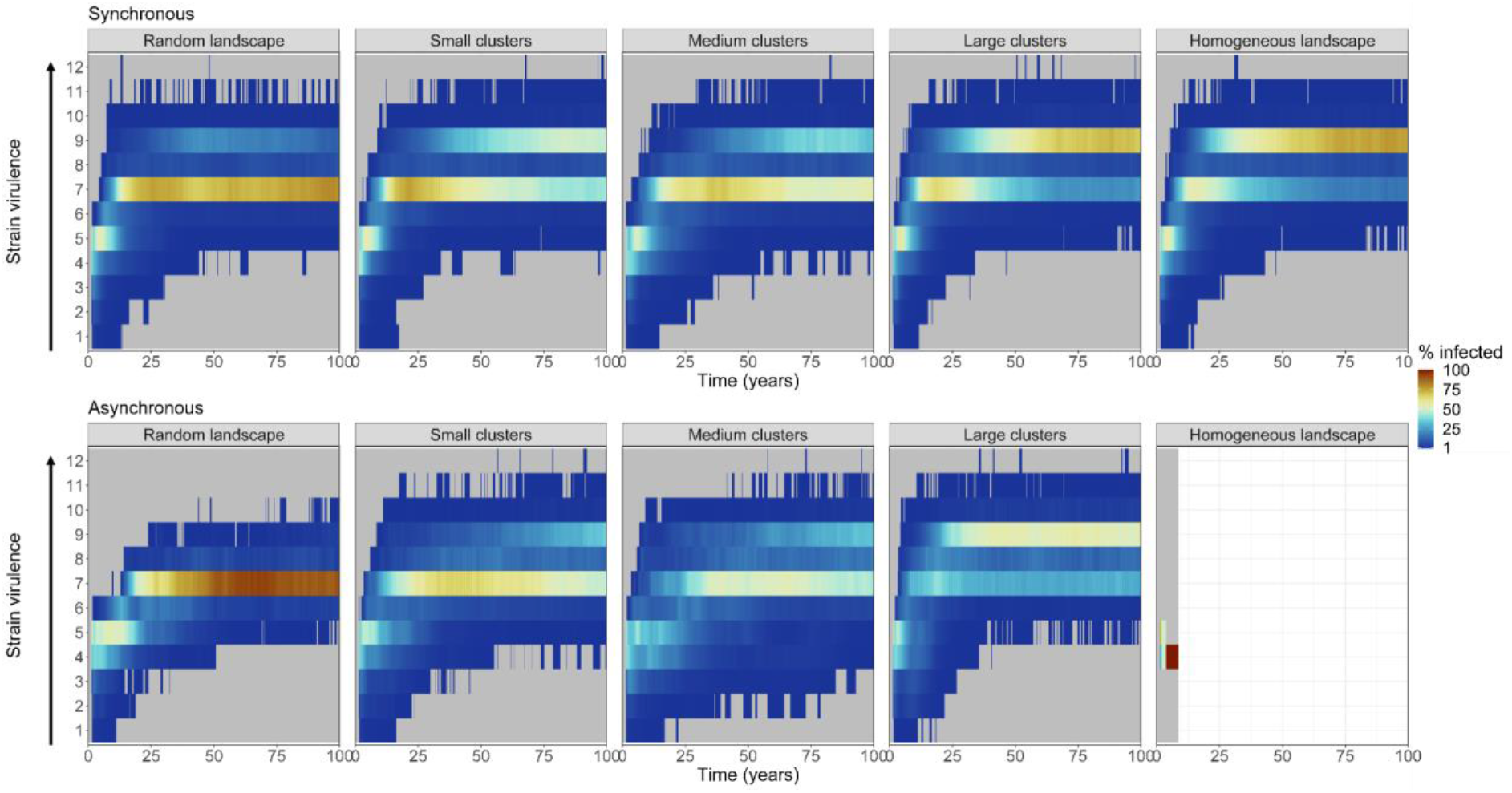
Occurrence and dominance of the different virulence strains in synchronous (t_lag_ = 0, top row) and asynchronous (t_lag_ = 100, bottom row) scenarios. Colour gradient represents the proportion of infected individuals with each strain in the landscape. Grey areas represent zero occurrence of the strains.

**Figure 4:**
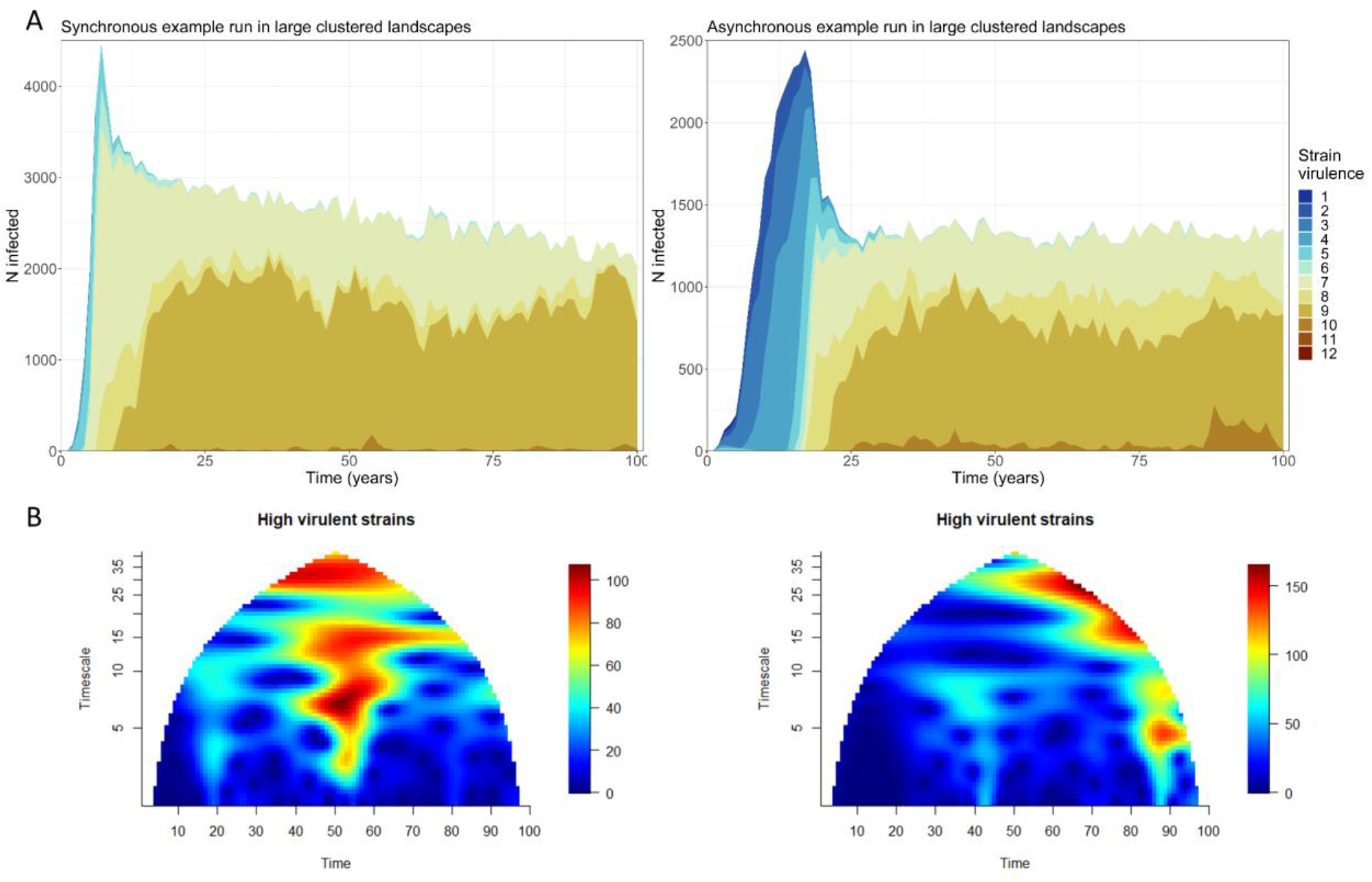
**A**-Muller plot for a single example run in a large clustered landscape in synchronous (left) and asynchronous (right) scenarios, showing the number of infected individuals for each strain (colour) over time aggregated as annual mean. **B**-Wavelet analysis of high virulent strains in synchronous (left) and asynchronous (right) scenarios of a single example run in a large clustered landscape using the R package “wsyn” [49].

## Discussion

To extend the understanding of pathogen evolution and spread during epidemics, we implemented virulence evolution in an individual-based model simulating an interdependent, tri-trophic system (landscape resources - host - pathogen) under the effects of global change. In accordance with our hypotheses, we found an increase in pathogenic virulence and a shift in strain dominance with increasing landscape homogenisation.

Landscape homogenisation alters the density distribution of susceptible host individuals by increasing host connectivity, which subsequently can lead to more infection events and viral mutations. The detrimental effect that density, connectivity and contact rates can have on viral mutations and infection events can also be observed in “superspreader”-events of the current SARS-CoV-2 pandemic (Tasakis et al. 2021). Our results support that host density and connectivity are the most important factors that affect the emergence of high virulence in directly transmitted diseases under classical transmission-virulence trade-offs (Castillo-Chavez and Velasco-Hernández 1998).While we found lower mean virulence in scenarios with asynchronous host resources, the landscape heterogeneity was the main driver of virulence evolution. Interestingly, under asynchrony, we found higher proportions of low and high strains coexisting in homogeneous landscapes, indicating that isolated disease hotspots (Kürschner et al. 2021) could facilitate the persistence of different viral strains.

As long as host populations in our model are distributed heterogeneously, mean pathogenic virulence remains similar, with little change from completely heterogeneous, i.e., random landscapes, to the less heterogeneous medium habitat clusters. However, in large clusters, a clear increase in mean virulence was apparent, showing that there is a threshold in landscape homogeneity not only enhancing disease spread, but also evolution towards higher virulence. These modelling findings are consistent with previous research on thresholds in disease transmission and functional connectivity. For example, homogenous landscapes have been shown to facilitate the spread of rabies in raccoons (*Procyon lotor*) (Brunker et al. 2012) or tuberculosis in badgers (*Meles meles*) (Acevedo et al. 2019), while more heterogeneous landscapes have been shown to limit the spread of highly virulent pathogens (Lane-deGraaf et al. 2013 p.). Host-pathogen interactions — in directly transmitted diseases — occur at specific locations and points in time, with the spatial and temporal variability in the availability of susceptible hosts being one of the governing factors of a successful transmission (Hudson 2002, Ostfeld et al. 2005, Real and Biek 2007). Consequently, homogenous landscapes and their lack of barriers allow more virulent pathogen strains to infect a sufficient number of hosts to persist in those landscapes. On the contrary, in heterogeneous landscapes, small clusters of high host density in a matrix of low density cause the extinction of highly virulent strains. This ‘dilution’ pattern can be explained by the short survival time of individuals in the matrix that form an immunity belt around the clusters and prevent spread between clusters (Marescot et al. 2021). Hence, in parallel with the ‘dilution hypotheses’ at the community scale, heterogeneous or ‘diverse’ landscapes provide less competent hosts for an epidemic (Patz et al. 2004, Civitello et al. 2015).

Increasing landscape homogenisation also resulted in higher mean virulence in scenarios with asynchrony between host life-history and resource availability (prediction 3). Even though overall susceptible host density was lower in asynchronous scenarios, the homogenous landscape’s increased connectivity allowed for higher virulent strains to persist at high prevalence. In the more homogenous, but still clustered, landscape, composed of large areas of high habitat suitability, the virulence of occurring strains was similar between the scenarios with and without synchrony. This indicates a strong effect of landscape configuration.

Interestingly, in our previous study (Kürschner et al. 2021), we showed that increasing spatial homogeneity of the landscape affected pathogen persistence negatively without pathogen virulence evolution. One reason behind this difference lies in the temporal differentiation of the strains within the landscapes. During the beginning of an outbreak, the pathogen strains with low virulence are able to spread across the landscape into larger habitat clusters due to the long survival times they impose on their hosts. Once the susceptible host density in one of the neighbouring areas is high enough, highly virulent strains that previously only occurred in low prevalence outcompete the low virulent strains and increase in prevalence. In other words, when host density increases, the high virulence strains capitalize on the high possibility for transmission and are likely to become dominant (Altizer et al. 2006, Hite and Cressler 2018). However, although highly virulent strains became more dominant, lower virulent strains continued to persist within the host population. In line with our findings, the coexistence of high and low virulent strains was also shown for rabbit haemorrhagic disease in the United Kingdom (Forrester et al. 2009) as well as influenza A in wild birds (Olsen et al. 2006).

Furthermore, our results show that, independent of landscape heterogeneity, a single, low virulent strain of a pathogen is able to evolve into a complex system of multiple coexisting strains with varying virulence (prediction 1). However, while multiple strains coexisted at any given time throughout all tested scenarios, we demonstrated that some strains likely become dominant (prediction 2). Similarly, a system of coexisting low and highly virulent strains were reported by empirical studies of the African swine fever virus in wild boar (Portugal et al. 2015), a pathogen causing severe diseases with huge economic impact (Artois et al. 2002). In this system the carriers of low virulent strains could remain infectious over long periods of time (de Carvalho Ferreira et al. 2012) increasing the chance of the pathogen transmission and its mutation into higher virulent strains, which could become dominant over time. In our study, the virulence of the dominant strain was intrinsically linked to the degree of landscape homogenisation but was also variable in time. Our findings are consistent with theoretical models that showed an increase of pathogenic virulence over time (Osnas et al. 2015). However, while Osnas et al. (2015) assumed a direct trade-off between virulence and host movement in homogenous landscapes, here we show that different landscape configurations may lead to the same patterns of increasing virulence without the necessity of such a trade-off.

On the one hand, our results show that with natural landscapes becoming more fragmented and resources becoming more asynchronous due to global change, a shift towards lower virulent pathogens could be expected. As a consequence, some diseases may become endemic in their respective host populations. The longer a pathogen is able to persist within its host population the higher the risk for spontaneous mutations and the possibility of spillovers to other species. On the other hand, global change will lead to increasing homogenisation within those fragments (Patz et al. 2004) and has the potential to increase the average pathogenic virulence with possibly catastrophic effects on wildlife communities. A large variance in virulence has been shown among infected host individuals, where the infection can range from severe to asymptomatic. This variation can be the result of a variety of factors, including genetic variation or intraspecific host interactions but also environmental conditions (Ebert and Bull 2003). Furthermore, an increase in virulence will go hand in hand with higher transmission rates in many diseases (Messinger and Ostling 2009, Alizon and Michalakis 2015) that will increase the probability of pathogen spillovers even more. While pathogen spillovers to other wild or domestic animal populations can have profound social or economic effects (Kamo et al. 2007), the possibly detrimental effects on human health cannot be underestimated. The current SARS-CoV-2 pandemic clearly highlights the importance of understanding which factors govern the spread of diseases in wildlife populations and how anthropogenic changes may alter those in the future.

## Methods

### Model overview

We modified a spatially explicit individual-based, eco-epidemiological model developed by Kürschner et al. (2021). It is based on earlier models considering neighbourhood infections only that was developed by Kramer-Schadt et al. (2009), Lange et al. (2012a, b) and Scherer et al. (2020) and includes spatiotemporal landscape dynamics representing changing resource availability, coupled with resource-based mortality. We incorporated evolution of viral traits such as virulence and corresponding trade-offs with viral transmission (see below). A complete and detailed model description following the ODD (Overview, Design concepts, Detail) protocol (Grimm et al. 2006, 2010) is provided in the supplementary material and the model (implementation) in the Zenodo Database and on GitHub [links provided on acceptance].

The model comprises three main components, a host model depending on underlying landscape features, an epidemiological pathogen model and a pathogen evolutionary model. Host individuals are characterised by sex, age, location, demographic status (residential, dispersing) and epidemiological status (susceptible, infected, immune). The epidemiological status of the individuals is defined by an SIR epidemiological classification (susceptible, infected, and recovered; Kermack and McKendrick 1927)). The pathogen is characterized by strain type, virulence and transmission. The pathogen model alters host survival rates and infection length depending on the pathogen’s virulence, while the dynamic landscape features determine host reproductive success. We record strain occurrences as the number of infected individuals carrying a specific strain and pathogen persistence, measured at the level of simulation runs (see below).

### Pathogen dynamics

We determined the course of the disease by an age-specific case fatality rate and a strain-specific infectious period. Highly virulent strains are characterized by a short infectious period and low virulent strains by a long infectious period. Transiently infected hosts shed the pathogen for one week and gain lifelong immunity (Dahle and Liess 1992). Infection dynamics emerge from multiple processes: within-group transmission and individual age-dependent courses of infection. Within groups, the density-dependent infection pressure (i.e. the chance of a host individual to become infected) is determined by a transmission chance and the number of infectious group members carrying the same strain. In this model we included the dependence of the transmission chance on the strain’s virulence, so that the strains with higher virulence have higher transmission chance. Furthermore, we modified the density dependence of the infection pressure to be strain-specific to accommodate a lower per-strain infection density for the following reason: The original model based on a single pathogen strain used the density of infected individuals in a cell to infer the likelihood for a susceptible host in that cell to become infected based on a binomial model.. Our model allows the evolution into 12 (arbitrarily categorized) different viral strains. The infection pressure λ, i.e. the probability of pathogen transmission to a susceptible host individual, is determined for each strain individually. Differences in strain transmissibility are added to the strain specific infection pressure through T_s_ (1). The probability *λ_is_* of an individual *i* of being infected by a specific strain *s* is calculated as

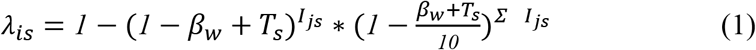

with *β_w_* being the individual probability of transmission to the power of all infected individuals *Ijs* in a group *j* per strain *s* as well as a reduced transmission probability between groups (i.e. cells) 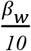 to the power of all infected individuals in neighboring groups *Σ I_js_*.

The strain virulence translates directly into infection length, i.e., host survival time, where a high virulence results in shorter survival times for the host compared to low-virulence. Consequently, the shorter lifetime of a highly virulent pathogen results in a shorter reproductive time span, while making the pathogen highly infective.

#### Evolution of pathogenic traits

Virulence and transmission are emergent properties and are evolving in the model. This means, while the position of each of the 12 strains on the transmission trade-off-curve is fixed, the selection of each strain during a transmission event is variable. Our trade-off curve is modelled to follow theoretical transmission-virulence trade-off curves (Alizon et al. 2009) and is applied for each infected host individually. During a transmission event, a strain can, with a mutation rate of 0.01, mutate into a new strain with a different virulence. The virulence of the new strain is selected from a normal distribution with a standard deviation σ = 1 around the virulence value of the originally transmitted strain, meaning that the new strain will be closely related to the parental strain.

### Landscape structure and dynamics

The tested landscapes consist of a spatial grid of 1.250 2 km x 2 km cells, each representing the average home range of a social host, e.g. a wild boar group (Kramer-Schadt et al. 2009), totalling a 100 km x 50 km landscape. The landscapes are self-contained systems without any outside interaction. Each cell is characterized by a variable resource availability that represents host breeding capacity and translates directly into host group size, with the minimum being one breeding female per group to a maximum of nine. Resource availability was adapted to achieve the average wild boar density of five breeding females per km^2^ (Howells and Edwards-Jones 1997, Sodeikat and Pohlmeyer 2003, Melis et al. 2006). We investigated several landscape scenarios of varying spatial complexity, ranging from a fully random landscape structure to different degrees of random landscape clusters generated in R (R Core Team 2020) using the NLMR package (Sciaini et al. 2018) up to a fully homogeneous landscape. To exclude any biases that could stem from different host densities, the mean female breeding capacity was kept constant at five females per km^2^ across the different landscape types (Supplementary material Appendix Fig. B1). The spatiotemporal landscape dynamics that were designed to mimic seasonal changes in resource availability by gradually increasing and decreasing resource availability were kept unchanged from the previous model implementation by Kürschner et al. (2021).

### Process overview and scheduling

The temporal resolution of the model equals the approximate pathogen incubation time of one week (Artois et al. 2002). The model procedures were scheduled each step in the following order: pathogen transmission, pathogen evolution, natal host group split of subadult males and females, resource-based host dispersal, host reproduction, baseline host mortality, strain-based host mortality, resource-based host mortality, host ageing and landscape dynamics. Natal group split of males and females was limited to week 17 and week 29 of each year, respectively, representing the observed dispersal time for each sex.

#### Host mortality

Mortality in response to resource availability remained unchanged to the previous model implementation (for details see ODD in the supplementary material). Additionally, we added a fixed, strain-specific mortality for each strain that affects the host population.

#### Landscape dynamics with temporal lag

We modelled two levels of temporal lag (t_lag_) implemented in Kürschner et al. (2021). We focus on the level 0% (synchrony between host population dynamics and resource availability) to 100% (asynchrony between host population dynamics and resource availability), with the latter simulating phenological mismatch between the resources and hosts reproduction potentially due to climate change. The extreme values were chosen because previous studies investigating temporal lag did not show strong effects in the intermediary steps (Kürschner et al. 2021).

### Model analysis

Each simulation was run for 100 years in total, with the virus released in a randomly taken week of the second year (week 53–104), to allow the population to stabilize after initialization. The virus was introduced to a set of multiple predefined cells in the centre of the landscape to ensure an outbreak. The virus was released in a low virulence variant. We ran 25 repetitions per combination of landscape scenarios (5 levels: small clusters, medium clusters, large clusters, homogenous landscape and random) and asynchrony (2 levels: t_lag_ 0%, t_lag_ 100%). We also analysed the strain occurrence (i.e., if a strain was present in any landscape cell, recorded at every timestep) and number of infected hosts per strain at every timestep to measure strain extinction as well as reappearance through mutation. We further recorded the proportion that each strain contributed to the pool of infected hosts by calculating the ratio of the hosts infected with each strain to the total number of hosts infected with all strains, at each time step. To highlight differences in strain composition in those scenarios, we subtracted the mean strain proportion in asynchronous scenarios from the mean proportion in synchronous scenarios. We categorized all viral strains into three categories: low virulence strains; medium virulence strains; high virulence strains, each compartment summing the outcomes of 4 of the 12 strains modelled.

## Acknowledgements

This work was supported by the German Research Foundation (DFG) in the framework of the BioMove Research Training Group (DFG-GRK 2118/1). We thank Volker Grimm for valuable comments on earlier drafts of this manuscript and Florian Jeltsch and Heribert Hofer for helpful discussions.

## Statement of authorship

All authors agree to submission of the manuscript, and each author carries a degree of responsibility for the accuracy, integrity and ethics of the manuscript and works described therein.

## Author contributions

TK and SKS developed the core idea and designed the study. TK rewrote and modified the simulation model together with CS and SKS. TK, VR, and SKS analysed the simulation results. TK is the lead author and CS, VR, NB and SKS contributed substantially to the writing. All authors agreed to submission of the manuscript, and each author is accountable for the aspects of the conducted work and ensures that questions related to the accuracy or integrity of any part of the work are appropriately investigated and resolved.

